# Hierarchical processing underpins competition in tactile perceptual bistability

**DOI:** 10.1101/2022.08.16.504072

**Authors:** Farzaneh Darki, Andrea Ferrario, James Rankin

## Abstract

Ambiguous sensory information can lead to spontaneous alternations between perceptual states, recently shown to extend to tactile perception. The authors recently proposed a simplified form of tactile rivalry which evokes two competing percepts for a fixed difference in input amplitudes across antiphase, pulsatile stimulation of the left and right fingers. This study addresses the need for a tactile rivalry model that captures the dynamics of perceptual alternations and that incorporates the structure of the somatosensory system. The model features hierarchical processing with two stages; a first stage resolves perceptual competition, leading to perceptual alternations; and a second stage encodes perceptual interpretations. The first stage could be located downstream of brainstem nuclei and the second stage could be located within the primary somatosensory cortex (area 3b). The model captures dynamical features specific to the tactile rivalry percepts and produces general characteristics of perceptual rivalry: input strength dependence of dominance times (Levelt’s proposition II), short-tailed skewness of dominance time distributions and the ratio of distribution moments. The presented modelling work leads to experimentally testable predictions. The same hierarchical model could generalise to account for percept formation, competition and alternations for bistable stimuli that involve pulsatile inputs from the visual and auditory domains.

**Author summary:** Perceptual ambiguity involving the touch sensation has seen increased recent interest. It provides interesting opportunity to explore how our perceptual experience is resolved by dynamic computations in the brain. We recently proposed a simple form of tactile rivalry where stimuli consisted of antiphase sequences of high and low intensity pulses delivered to the right and left index fingers. The stimulus can be perceived as either one simultaneous pattern of vibration on both hands, or as a pattern of vibrations that jumps from one hand to the other, giving a sensation of apparent movement. During long presentation of the stimuli, one’s perception switches every 5–20 seconds between these two interpretations, a phenomenon called tactile perceptual bistability. This study presents the first computational model for tactile bistability and is based on the structure of sensory brain areas. The model captures important characteristics of perceptual interpretations for tactile rivalry. We offer predictions in terms of how left-right tactile intensity differences are encoded and propose a location for the encoding of perceptual interpretations in sensory brain areas. The model provides a generalisable framework that can make useful predictions for future behavioural experiments with tactile and other types of stimuli.

## Introduction

Perceptual rivalry occurs when sensory information is ambiguous and the brain cannot commit to a single percept; instead, it switches between mutually exclusive interpretations every few seconds [1]. Examples of perceptual rivalry span across different sensory modalities including vision [2–5], audition [6] and olfaction [7]. In the tactile domain, perceptual rivalry was introduced with a tactile illusion based on the visual apparent-motion quartet [8–11]. Recent experiments with vibrotactile stimuli have shown that several of the general characteristics of perceptual rivalry extend to tactile domain [12]. Vibrotactile stimuli consisted of antiphase sequences of high and low intensity high frequency pulses delivered to the right and left index fingers (Fig 1A. Participants perceived the stimulus (Fig 1B) as either one simultaneous pattern of vibration on both hands (SIM), or patterns of vibration that jumped from one hand to the other hand, giving a sensation of apparent movement (AM) [2], and for long presentations of the stimulus (*>* 30 *s*), perception switched back and forth between these two perceptual interpretations (percepts).

**Fig 1.**
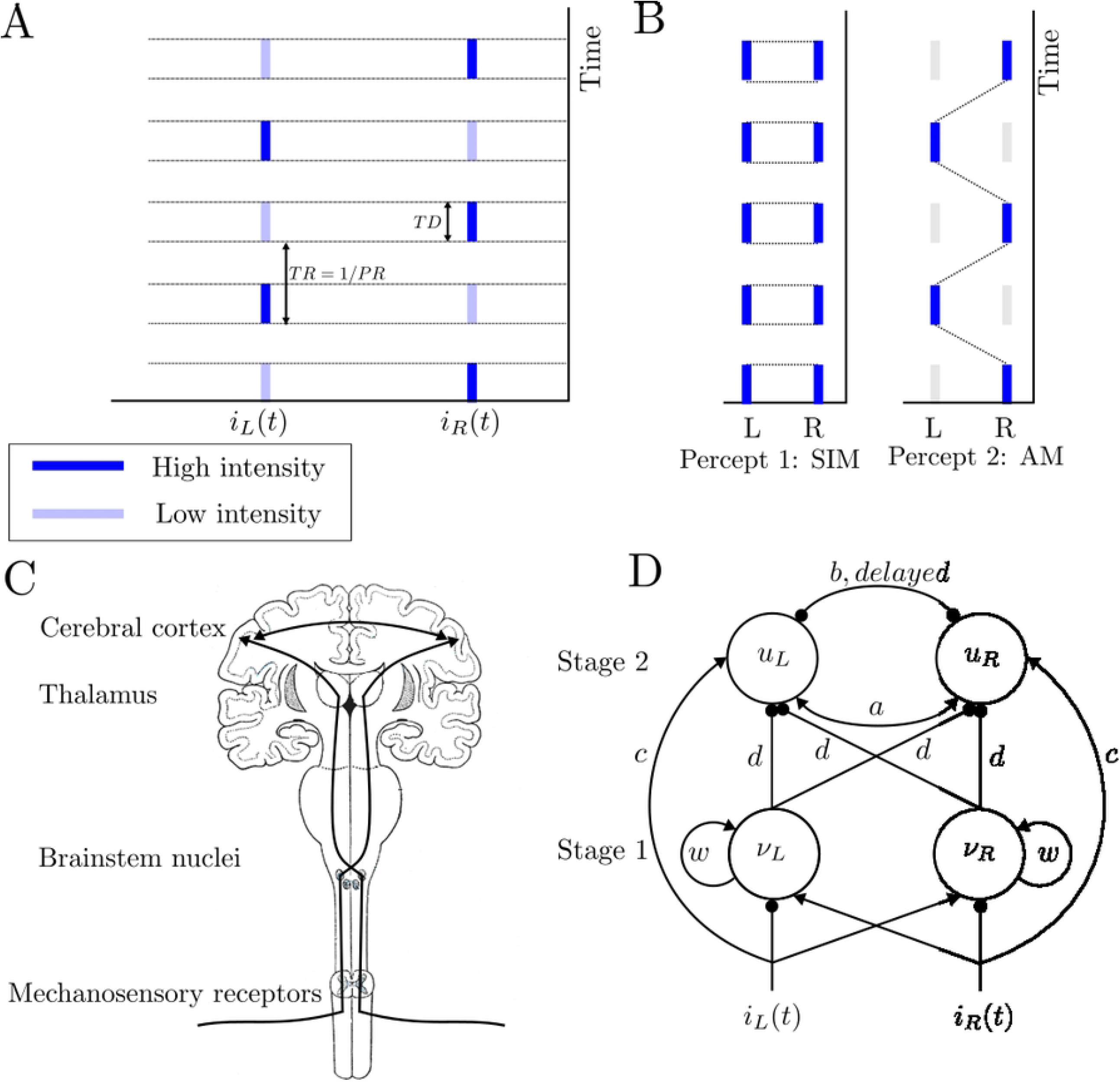
**(A) Vibrotactile stimuli.** Vibrotactile stimuli consist of antiphase sequences of high (dark blue) and low-intensity (light blue) 400 ms duration, 200 Hz pulses delivered to the right and left index finger each followed by a 400 ms gap, i.e. the pulse duration *TD* = 0.4 s and pulse repetition time *TR* = 0.8 s. **(B) Percept types**. During a trial, the participant’s perceptual interpretation of the stimuli changes. When the patterns are played with equal intensity, they are unambiguously perceived as one simultaneous vibration (SIM). With a fixed intensity difference (Δ*I >* 0 *dB*) between the high- and low-intensity tactile pulses, perception switches back and forth between two percepts: SIM (perceived as a fixed intensity on each hand, even though the intensity is changing) and AM (perceived as pulses of vibrations jumping from one hand to the other hand). **(C) Structure of somatosensory pathway**. Afferent fibres cross over and project to thalamic nuclei on opposite side, then project to cerebral cortex. **(D) Schematic of the model of tactile rivalry**. Inhibitory connections are shown with filled circles, and excitatory connections with black arrows; see text for definitions of neural populations (units) and their parameters.

There are numerous stimulus examples for perceptual ambiguity across different sensory modalities and across different paradigms within the same sensory modality. In spite of this diversity, general characteristics of this phenomenon appear to be quasiuniversal. Firstly, Levelt’s propositions have been widely used to describe perceptual rivalry in the visual [13, 14], auditory [15] and recently in tactile domains [12]. For example, the generalization of Levelt’s proposition II states that increasing the difference between percept strengths increases the mean perceptual dominance of the stronger percepts [16]. Secondly, despite mean dominance times varying widely in multistable experiments, across different observers and stimulus contrasts [17, 18], the statistical distribution of perceptual phases maintains a constant shape, resembling a log-normal or gamma distribution with consistent values for the coefficient of variation *cv* and skewness ratio (*γ*_1_*/cv*; where *γ*_1_ is the skewness) [12, 19, 20]. Thirdly, the dominance durations of successive percepts are correlated positively for perceptual phases that were one phase apart (between different percepts) [12, 21–23]. These similar properties in multistable phenomena suggest that the underlying mechanisms may be general.

Computational models of perceptual alternations have helped significantly with our understanding of sensory processing across visual and auditory domains. These models focus on the neural processing of sensory information, the dependence of average switching times on stimulus parameters and the statistical distribution of times between switches [15, 18, 24–29]. Perceptual bistability results from competition between units representing neural populations associated with different percepts (e.g. units driven by inputs from the left and right eyes in binocular rivalry) [23, 26, 29]. General models of rivalry usually incorporate a slow process together with reciprocal inhibition to produce perceptual alternations. Bifurcation analysis is utilized to compute different dynamical regimes and boundaries between them for multiple parameters with fixed [25] and periodic stimuli [30].

Here, we address the need for a tactile rivalry model (to the best of our knowledge, one is yet to be proposed) that accounts for well-established results on the duration of dominance intervals and incorporates the structure of the somatosensory system based on physiological evidence. It remains an open modelling challenge to address how sequences of pulsatile inputs are encoded as percepts, and how neural competition resolves ambiguity to select and switch between percepts. Existing models with pulsatile inputs did not address the competition mechanisms that link between inputs separated in time and across a feature (e.g. between left and right eyes in binocular rivalry or across a tone frequency in auditory streaming) [15, 26, 29]. However, a recent study (upon which we build) went some way to addressing the question of percept formation [31].

In this study, we develop a mathematical model of tactile rivalry that focuses on accurately reproducing the dynamics of the perceptual alternations. The model is neuromechanistic, i.e. based on computational principles widely accepted as underpinning cortical processing. This formulation is directly motivated from physiological studies of tactile perception (as reviewed below) [32–34], a model of bistable dynamics [35], and a model of auditory percept encoding for sequences of tones [31]. The model of tactile rivalry presented here consists of two processing stages; first stage for producing perceptual alternations; and a second stage for encoding the percept types (SIM and AM). The powerful combination of bifurcation analysis along with optimisation tools have been used to tune certain features of the model.

The model presented here is able to produce experimentally-observed characteristics of tactile rivalry, including the specific temporal structure of the percepts. We offer predictions in terms of how left-right tactile intensity differences are encoded and the putative location of percept encoding in somatosensory cortex. The model provides a framework to predict parameter dependence of dominance duration for future behavioural work. The same framework and hierarchical structure can generalise to the study of other perceptually bistable stimuli involving sequences of pulsatile inputs as investigated elsewhere in auditory [15] and visual paradigms [26, 29].

## Materials and methods

### Background on somatosenory pathways

We give a brief overview of literature on the neural encoding of vibrotactile features in the somatosensory system. The spatio-temporal characteristics of tactile perception have been investigated with a range of experimental approaches. At the somatosensory periphery, the firing rate response of tactile fibres to a vibration is dependent not only on its amplitude but also on its frequency. Therefore, the amplitude information carried in the firing rates of any population of tactile fibres is ambiguous. Rather, the intensity of skin vibrations is encoded in the firing rate evoked in all tactile nerve fibres, weighted by fibre type [36]. So it is unclear how these two stimulus dimensions can be independently decoded by downstream structures [37].

The different types of tactile receptors differ in their frequency sensitivity profiles. Slowly adapting fibres (SA1) tend to be more sensitive at low frequencies (below about 10 Hz), PC fibres (RA2) peak in sensitivity at around 250 Hz (respond in range 40-400 Hz), and rapidly adapting fibers (RA1) prefer intermediate frequencies (5-50 Hz) [38]. Neural signals from a single class of receptors can convey ambiguous information. Afferent signals from different types of mechanoreceptors are combined to give rise to tactile percepts. A striking aspect of afferent responses to skin vibrations is that they exhibit phase locked response to periodic vibratory stimuli (that is, they produce one spike or burst of spikes within a restricted portion of each stimulus cycle) as do their counterparts in the auditory nerve [39, 40]. This temporal patterning in afferent responses carries information about the frequency composition of vibratory stimuli applied to the skin [41]. However, how afferent spiking activity translates into the perception of frequency is still unknown. Neurons in somatosensory cortex also exhibit phase locking to low-frequency vibrations, but the constancy of the phase locking decreases rapidly as vibratory frequency increases.

### Percepts and model inputs

During experimental trials of tactile rivalry, stimuli presented in Fig 1A are perceived as either one simultaneous pattern of vibration on both hands (SIM), or patterns of vibration that jumped from one hand to the other hand, giving a sensation of apparent movement (AM) (Fig 1B).

In our tactile rivalry experiments [12], stimuli consisted of sequences of high (H) and low (L) intensity vibratory pulses, each followed by a silent interval (antiphase sequences of *H* – *L* − *H* − *L* for the right hand and *L*− *H* – *L* − *H* for the left hand, “− “ indicates the silent gap) (Fig 1A). The intensity of the *L* stimulus is Δ*I* below the intensity of the H stimulus on a logarithmic scale (*dB*). Model input functions, *i*_*R*_(*t*) and *i*_*L*_(*t*) as defined by Eq (1) and (2), are antiphase periodic square waves [31] corresponding to the stimuli delivered to right and left hands, respectively.

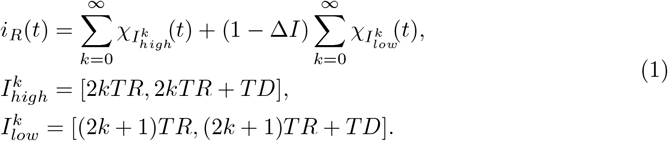

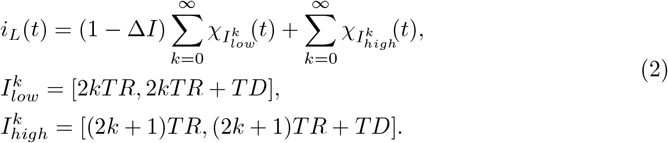

Where Δ*I* represents the intensity difference between high and low amplitude vibratory pulses. *χ*_*I*_ is the standard indicator function over the set of intervals *I*, defined as *χ*_*I*_ (*t*) = 1 for *t* in *I* and 0 otherwise. The intervals when high and low intensity vibrations are on are respectively given by 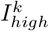 and 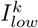. The parameter *TD* represents the duration of high or low intensity pulses, and *TR* is the time between pulse onsets (*PR* = 1*/TR* is the presentation rate); see a schematic plot of the stimulus in Fig 1A and refer forward to a plot of one period of the stimulus in S3 FigA.

In order to have a smooth square waveform rather than an ideal discontinuous square waveform, we used a steep sigmoid,

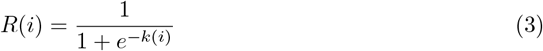

with k=20 which defines the slope. So instead of the *i*_*R*_ and *i*_*L*_, we substitute *R*(*i*_*R*_) and *R*(*i*_*L*_) in the model’s inputs.

### Processing stages

Here we want to introduce a computational model of tactile rivalry to capture the characteristics of this phenomenon as observed in the experiments. To this end, we consider how inputs from the left and right hands project to primary somatosensory cortex (S1) and how features like amplitude, frequency and timing are encoded there (Fig 1C). The model presented here consists of two processing stages (Fig 1D); a first stage with units *ν*_*L*_ and *ν*_*R*_ generates perceptual alternations, and a second stage with units *u*_*L*_ and *u*_*R*_ encodes the percepts (SIM and AM). The two stages receive the inputs defined above in parallel. The second stage encodes the temporal pattern associated with each percept directly from the stimulus based on mutual fast excitation slow and delayed inhibition as described in [31]. The first stage computes amplitude differences between the left and right inputs (as justified below) and amplifies these differences via global inhibition of the second stage (strength *d*); however, this effect is transient due to adaptation (a slow negative recurrent feedback) on each unit *ν*_*L*_ and *ν*_*R*_, which leads to alternations. As described in more detail below, alternations in the first stage lead to switches in the percept encoded by the second stage due to changes in the strength of global inhibition via *d*.

In addition to contra-lateral excitation, ipsi-lateral inhibition has been observed during unilateral touch in S1 [42, 43]. These effects are shown in Fig 1D, as inputs *i*_*R*_ and *i*_*L*_ have an excitatory effect on the opposite side (excitation of units *ν*_*L*_ and *ν*_*R*_, respectively), and inhibitory effects on the same sides (inhibit units *ν*_*R*_ and *ν*_*L*_, respectively). Thus, units *ν*_*R*_ and *ν*_*L*_ in the first stage effectively receive *i*_*L*_ − *i*_*R*_, and *i*_*R*_ − *i*_*L*_, respectively, which are antiphase pulses with amplitude equal to Δ*I* (Fig 2A).

**Fig 2.**
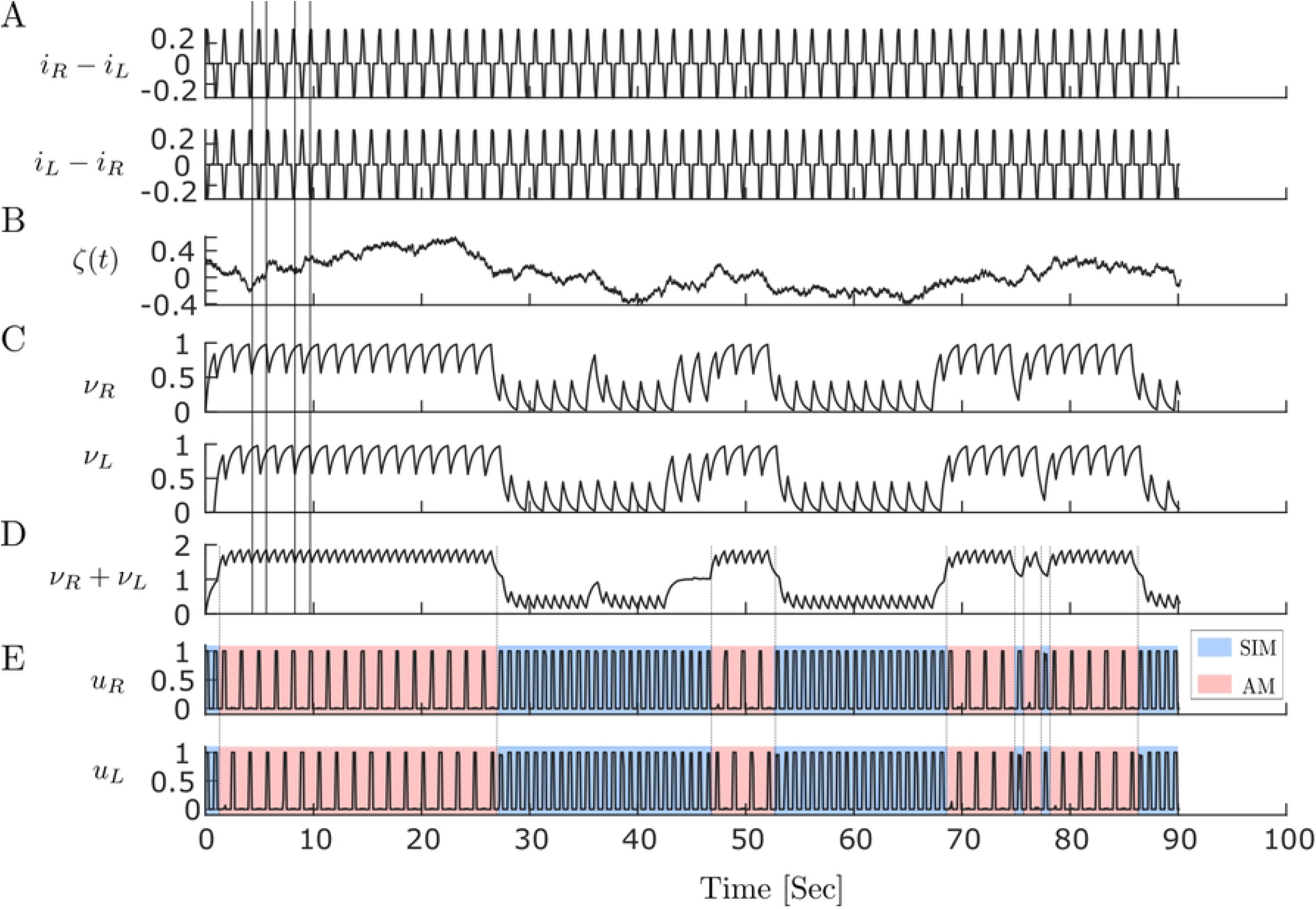
Time histories of tactile rivalry model. Population firing time responses at Δ*I* = 2 *dB* for 90 s simulation. **(A)** Net inputs to the left (top panel) and right (bottom units of the first stage). **(B)** Noise with parameters *σ* = 0.3 and *τ*_*n*_ = 0.05 which is added to the first stage inputs. **(C)** Firing activities of the first stage units to the inputs in panel A and noise in panel B. Vertical lines are plotted to show that unit activities in the first stage are antiphase due to their antiphase inputs. The activity decays when the input is negative and increases otherwise. **(D)** Sum of firing activities of the first stage units, which will be the inhibitory input to the both units of second stage. As mean firing activities in the first stage are antiphase, there is low amplitude oscillation when they are summed up. **(E)** Firing activities of the second stage units to the inputs in panel D. Perceptual alternation between SIM and AM percepts are seen as transition occurs between DOWN and UP states in panel D

The units *ν*_*R*_ and *ν*_*L*_ in the first stage are based on the simplified firing rate model from [35] with recurrent excitation and slower recurrent adaptation. These units produce alternations between an UP (ON) and DOWN (OFF) state. This model, repurposed as a component in the present study has previously been used to investigate the state of hippocampal and neocortical populations during NREM sleep (UP/DOWN states observed as spontaneous transitions during sleep). The analysis in their study provides a useful reference to tune model parameters. When the first stage units are in the DOWN state, inputs drive the second stage in its default setting where typically SIM is encoded, unless Δ*I* is very large. If the first stage units are in the UP state, inputs driving the second stage are less excitatory, leading to AM, unless Δ*I* is very small. The dynamics of the units model in the first stage are further described in S1 Appendix.

The second stage in Fig 1D shows units *u*_*R*_ and *u*_*L*_ that receive direct ipsi-lateral excitatory inputs from the right and the left side (*i*_*R*_ and *i*_*L*_, respectively), and also inhibitory ipsi- and contra-lateral connections through the earlier stage (units *ν*_*R*_ and *ν*_*L*_). Existence of these direct ipsi-lateral excitation is motivated by the fact that some neurons in S1 display bilateral hand receptive fields [34, 44]. Interhemispheric connections at the second stage are assumed to exist between units *u*_*R*_ and *u*_*L*_ through direct fast excitation with strength *a*, and the delayed, slowly decaying inhibition with strength *b*. Dynamics of units *u*_*R*_ and *u*_*L*_ are described in more details in S1 Appendix.

### Full tactile rivalry model with noise

To form the model of tactile rivalry, the model encoding alternations at the first stage is incorporated with the model encoding percepts at the second stage. Units *ν*_*R*_ and *ν*_*L*_ in the first stage make inter- and intra-hemispheric inhibitory connections with the units *u*_*R*_ and *u*_*L*_ in the second stage (Fig 1D).

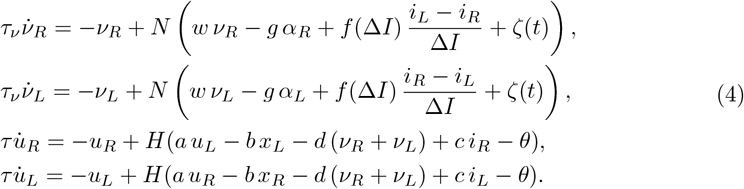

The dynamics of units in the first stage are described in terms of the mean firing rates *ν*_*R*_ and *ν*_*L*_ with time scale *τ*_*ν*_, and activity-driven adaptation *α*_*R*_ and *α*_*L*_ with time scale *τ*_*α*_ (Fig 1D). Where *w* is the strength of recurrent excitation, *g* is the strength of adaptation, (*i*_*L*_ − *i*_*R*_) and (*i*_*R*_ − *i*_*L*_) are the stimulus differences from the right and left, and *ζ*(*t*) is noisy fluctuations. The nonlinear function *f* (Δ*I*) will be determined later with data-driven optimisation. *N* (*x*) is assumed to be sigmoidal activation function as follows;

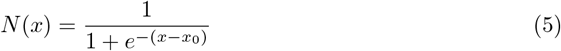

The dynamics of units in the second stage are described in terms of the mean firing rates *u*_*R*_ and *u*_*L*_ of two neural populations which encode sequences of vibratory input pulses with timescale *τ*. The synaptic variables *s*_*R*_ and *s*_*L*_ describe the time-evolution of inhibitory dynamics. Indirect synapses are also modelled by two other synaptic variables *x*_*R*_(*t*) and *x*_*L*_(*t*) that can generate delays approximately equal to *δ* [45]. The Heaviside gain function *H*(*x*) is equal to 1 for *x* > = 0, and 0 for *x* < 0. Mutual coupling through direct fast excitation has strength *a*. The delayed, slowly decaying inhibition has timescale *τ*_*i*_, strength *b* and delay *δ*. The strength of inhibitory connections between the first and second stage is *d*. The model is driven by excitatory inputs *i*_*R*_(*t*) and *i*_*L*_(*t*) with strength *c*. Here, *θ*_*u*_ and *θ*_*s*_ are activation thresholds, *α*_*i*_ and *β*_*i*_ (*i* = *s, x*) are positive constants.

We extended the previous model proposed by Ferrario and Rankin [31] by transforming the delayed inhibition into a system of ordinary differential equations using the approach described in [45]. This in turn allowed us to track periodic orbits modulated by forcing under parameter variation (as described in [30] and using numerical continuation with Auto07p). Note that this approach only works for small to moderate delays.

The Heaviside function *H*(*x*) that appears in the right-hand side of the full tactile model is a discontinuous function in its first derivative. Numerical continuation routines require smooth systems of equations. In order to solve this problem we have used a steep sigmoid function to smooth out the transition at zero.

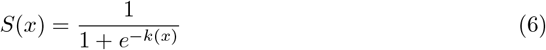

with k=20 which defines the slope. So instead of the *H*(*x*), we substitute *S*(*x*) in the right-hand side of the full tactile model.

Noise *ζ*(*t*) was implemented using Ornstein-Uhlenbeck model described by

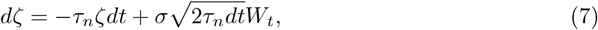

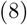

where *W*_*t*_ is a Wiener process with time scale *τ*_*n*_ and standard deviation *σ*.

### Bifurcation and statistical analysis

Bifurcation analysis of the model in the absence of noise was carried out with Auto07p [Source code for the model is available in the GitHub repository farzaneh-darki/Darki2022-hierarchical: https://github.com/farzaneh-darki/Darki2022-hierarchical]. For the statistical analysis of dominance duration distributions, the same model was implemented in MATLAB for simulations with noise. Numerical integration of the resulting stochastic differential equation was carried out using a standard Euler-Muruyama scheme with time step 0.01 ms which is much smaller than the fastest timescale (*τ* = 0.001 s=1 ms). All the model parameters and their corresponding values are provided in Table 1.

**Table 1.** Parameters of the model with their corresponding values used in the simulations. The main value of the model parameters are provided here. If a parameter changes from its main value, the new value determined in the relevant figure.

| Parameters | Values | Parameters | Values |
| --- | --- | --- | --- |
| $w$ | 6 | $a$ | 3.4 |
| $g$ | 1.5 | $b$ | 2.8 |
| $\tau_\nu$ | 0.9 | $c$ | 5.5 |
| $\tau_\alpha$ | 4.5 | $\delta$ | 0.005 |
| $x_0$ | 5 | $\tau_i$ | 0.25 |
| $k_A$ | 15 | $\tau$ | 0.001 |
| $\nu_0$ | 0.5 | $\theta$ | 0.5 |
| $\sigma$ | 1 | $\alpha_x$ | 50 |
| $\tau_n$ | 0.05 | $\beta_x$ | 8 |
| $d$ | 2.6 | $\theta_s$ | 0.22 |

### Simplified tactile rivalry model

As the inputs (*i*_*L*_ − *i*_*R*_) and (*i*_*R*_ − *i*_*L*_) are antiphase and there is also symmetry in the first stage of the tactile rivalry model, this model can be simplified, and units *ν*_*R*_ and *ν*_*L*_ can be replaced by one adapting recurrent model with variables *ν* and *α* and input *D* = *f* (Δ*I*) (Fig 3A). Where Δ*I* is a positive constant, and *f* is a nonlinearity to be determined by data-driven optimisation. So the simplified tactile rivalry model is described by

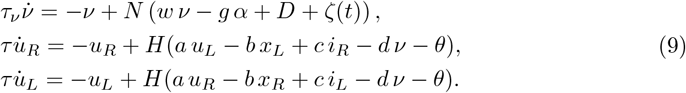

**Fig 3.**
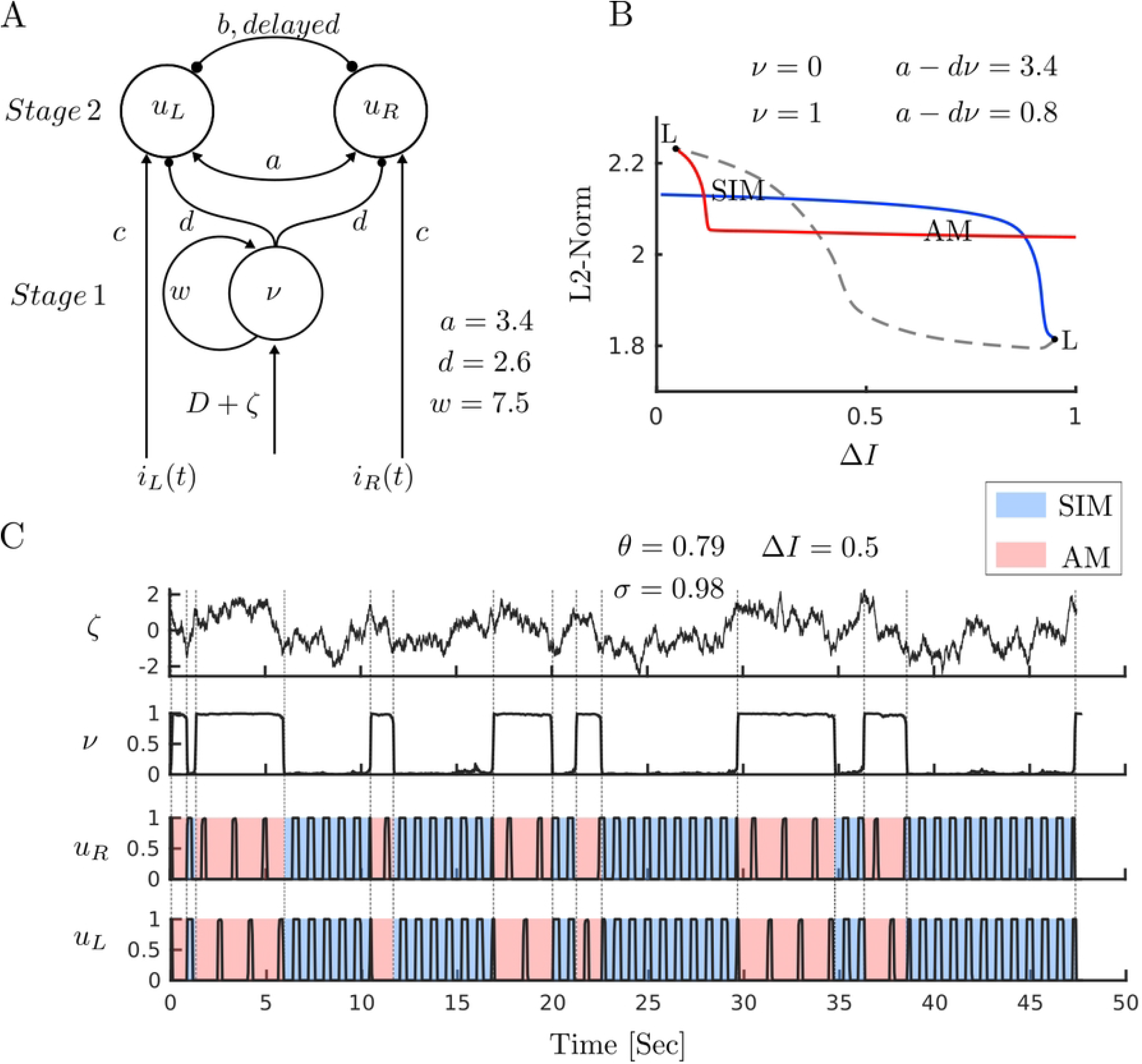
Mechanism of perceptual alternations. **(A) Simplified model of tactile rivalry.** The adapting recurrent model with firing rate *ν* (analysed in isolation in S1 Fig) makes inhibitory connections with strength *d* to the units encoding the percepts *u*_*R*_ and *u*_*L*_ (analysed in isolation in S3 Fig). **(B) Bifurcation analysis with respect to intensity difference** Δ*I*. There is a region of bistability between two fold of limit cycle bifurcation points (L). Branches of periodic orbits associated with SIM and AM percepts coexist at this interval. **(C) Time histories of model responses**. Population activities are simulated for 50 s at Δ*I* = 0.5 (*D* = 1.25). Noise realization (top panel), UP/DOWN alternations of the first stage unit driven by noise (second panel, *g* = 0), firing activities of the second stage unit (two bottom panels). Perceptual switching times are shown between SIM (blue) and AM (red) with dashed lines.

### Bistability in the simplified model

Analysis of the model’s component stages provided a means to tune parameters of the more tractable the simplified model, which could then be used in the full model to capture the desired dynamics, as will be discussed in the results section. Full details of this analysis of each stage is given in S1 Appendix. The general aim was to tune parameters so as to produce a region of bistability between states representing SIM and AM in the noise-free simplified model. The introduction of noise can then drive alternations between these states. Fig 3B shows a bifurcation diagram with the desired coexistence of SIM and AM solution branches over a significant range of Δ*I*. Fig 3C shows a time history of the noise-driven simplified model, where alternations in stage 1 (as *ν* transition occurs from zero to one or vice versa) drive switches between a synchronised SIM state (blue) and antiphase oscillations (AM). Population activities are simulated for 50 s at Δ*I* = 0.5. The top panel shows the noise realization with parameters *σ* = 0.79 and *τ*_*n*_ = 0.98.

For the interested reader, a brief summary of the bifurcation analysis of the component stages leading to Fig 3B is discussed here, with full details given in S1 Appendix. The adapting recurrent model in the first stage makes inhibitory connections with strength *d* to the model encoding the percepts in the second stage (Fig 3A). As seen in (S1 FigB & 6D) unit *ν* has a region of bistability where *ν* can be either zero or one (when input *D* lies between two fold bifurcation points). As this unit inhibits *u*_*L*_ and *u*_*R*_ with strength *d*, the convergence to 0 or 1 of this unit will modify the excitatory net inputs to units *u*_*L*_ and *u*_*R*_ and thus shift the branch of periodic orbits in Fig 3B to the right and to the left (when *ν* = 1, second stage receives less excitation, effectively *a*_eff_ = *a* − *dν*; see S3 FigE). This results in bistability between SIM and AM dynamical states shown in the bifurcation diagram of the whole model (Fig 3B).

## Results

Here we go through a qualitative description of the dynamics produced by the full tactile rivalry model presented in Eq (4) and explain how this qualitatively matches perceptual interpretations and alternations observed in tactile rivalry experiment. We further analyse the dependence of mean dominance durations and their variability (as characterised by a skewed distribution) on the stimulus parameter Δ*I*.

### Time history simulations of full tactile rivalry model

We first discuss the output from individual simulations of the model and illustrate how model’s firing rate variables can encode the competing percepts and perceptual alternations. A region of bistability, identified by a detailed bifurcation analysis of the model, was described in methods section above (Fig 3). For the interested reader, a detailed analysis of the model and it’s component stages as given in S1 Appendix shows how the tactile rivalry model was designed to encode percepts and generates perceptual alternations.

A 90 s time simulation for the full tactile rivalry model is shown in Fig 2. The units in the first stage are excited by the contra-lateral stimulus and inhibited by the ipsi-lateral stimulus. Thus, the net inputs to the left and right units of the first stage will be the contra-lateral stimuli minus ipsi-lateral stimuli. These inputs are antiphase pulses with amplitude proportional to Δ*I* as shown in Fig 2A. These inputs weighted by *f* (Δ*I*) and delivered to the units in the first stage. Noise is added to these inputs with amplitude *σ* = 0.3 and timescale *τ*_*n*_ = 0.05 (Fig 2B). Firing activities of these units in the first stage in response to the stimuli and noise are shown in Fig 2C. In the absence of noise, these adapting recurrent units could oscillate between the UP and DOWN states regularly. However, these oscillations are now driven by both adaptation and noise process. Fig 2D shows the sum of firing activities of the first stage, which is delivered as an inhibitory input to the both units of second stage. Firing activities of the second stage units to these inputs are shown in Fig 2E. Units of the second stage encode the SIM percept (both units fully respond to the high and low intensity pulses in the inputs) when there is low levels of inhibition from the first stage. When the level of inhibition crosses a certain threshold, the units in the second stage encode AM percept (both units only fully respond to the high intensity pulses in the inputs). Perceptual alternation between the SIM and AM percepts are seen as transitions between DOWN and UP states occur in the inhibitory inputs (Fig 2D&E).

### Stimulus parameter dependence

Results from experiments with vibrotactile stimuli demonstrate that Levelt’s proposition II holds in tactile domain [12]. Increasing intensity difference, causes the mean dominance of SIM percept to decrease and AM percept to increase. Mean dominance duration for both the perceptual durations from the model and the experiment (Experimental data from [12]) are plotted against intensity difference (Δ*I*) in Fig 4. The parameters of the noise (*σ, τ*_*n*_), time constant of adaptation (*τ*_*a*_), and nonlinearity in the inputs of the first stage (*D* = *f* (Δ*I*)) were determined using a genetic algorithm. Our optimisation approach also determined the nonlinearity *f*. The good match with experimental data, obtained by tuning a small number of parameters and the input nonlinearity offers confidence that the model presented here is an effective, parsimonious description of the potential mechanisms driving tactile rivalry.

**Fig 4.**
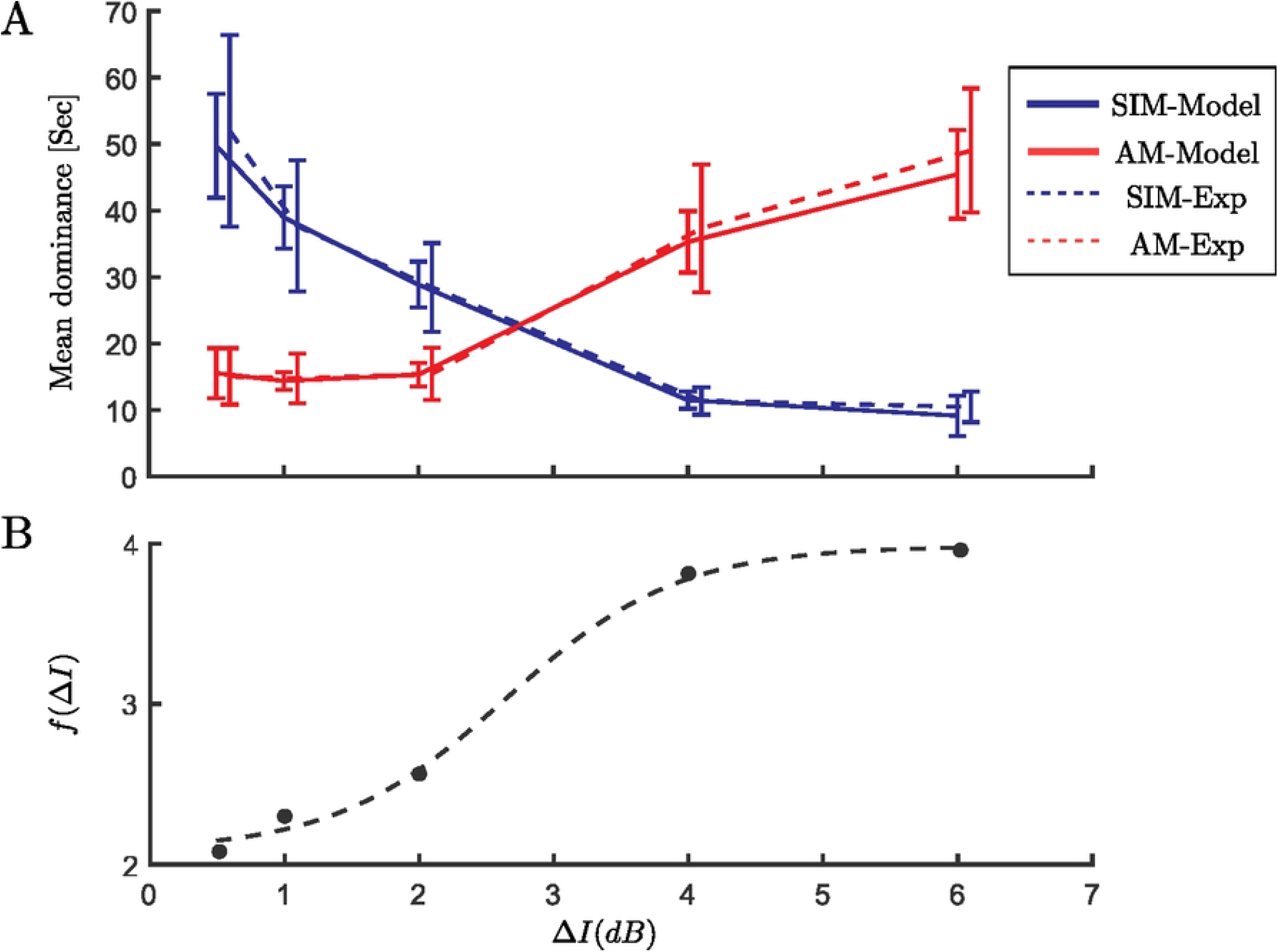
Levelt’s proposition II. **(A)** Experimental data are dashed curves and computational data from the model are solid curves with data points at different values of intensity difference Δ*I* = 0.5, 1, 2, 4, 6 on the x-axis, error bars show standard error of the mean. Mean dominance duration of the SIM percept (blue) increases as the intensity difference increases, while an opposite effect is observed for the AM percept (red). **(B)** Nonlinearity in the inputs of the first stage (*D* = *f* (Δ*I*)) are determined using an optimisation algorithm. Dashed black curve is the best fit for an offset and scaled sigmoid nonlinearity.

To find nonlinear function *f* (Δ*I*), we first estimated some points of it at the experimental conditions (Δ*I* = 0.5, 1, 2, 4, 6 *dB*) using a genetic algorithm (*f* (0.5) = 2.16, *f* (1) = 2.3, *f* (2) = 2.6, *f* (4) = 3.75, *f* (6) = 3.92 with *τ*_*n*_ = 0.05, *σ* = 0.3, *τ*_*a*_ = 5 *s*). Having these points, we showed an offset and scaled sigmoid function like;

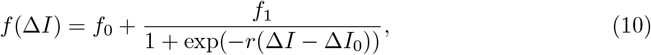

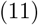

with parameters: *f*_0_ = 2.12 (offset), *f*_1_ = 1.80 (scale), *r* = 1.57 (slope), Δ*I*_0_ = 2.64 (equidominance), fits best to these points.

### Variability of perceptual durations

The distributions of normalized perceptual durations from the model and from the experiment are shown in Fig 5A&B. These distributions were compared with the gamma and log-normal distributions using a one-way Kolmogorov-Smirnov (KS) test. The null hypothesis is that the test data are drawn from the standard comparison distribution and a significant result (*p* < .05) indicates that the test data are not drawn from the comparison distribution. The one-way KS tests shows that the results produced by the tactile rivalry model best fit by a log-normal distribution, but that the gamma distribution can be rejected (*p*(*gamma*) < .05). For the experimental data, neither distribution could be rejected. However, in similar experiments with auditory bistability [15, 20] and visual bistability [20] a log-normal distribution provided a better fit than the gamma distribution. We suspect that increasing the number of participants, or number of trial repetitions, may offer a more conclusive result for tactile rivalry in the future.

**Fig 5.**
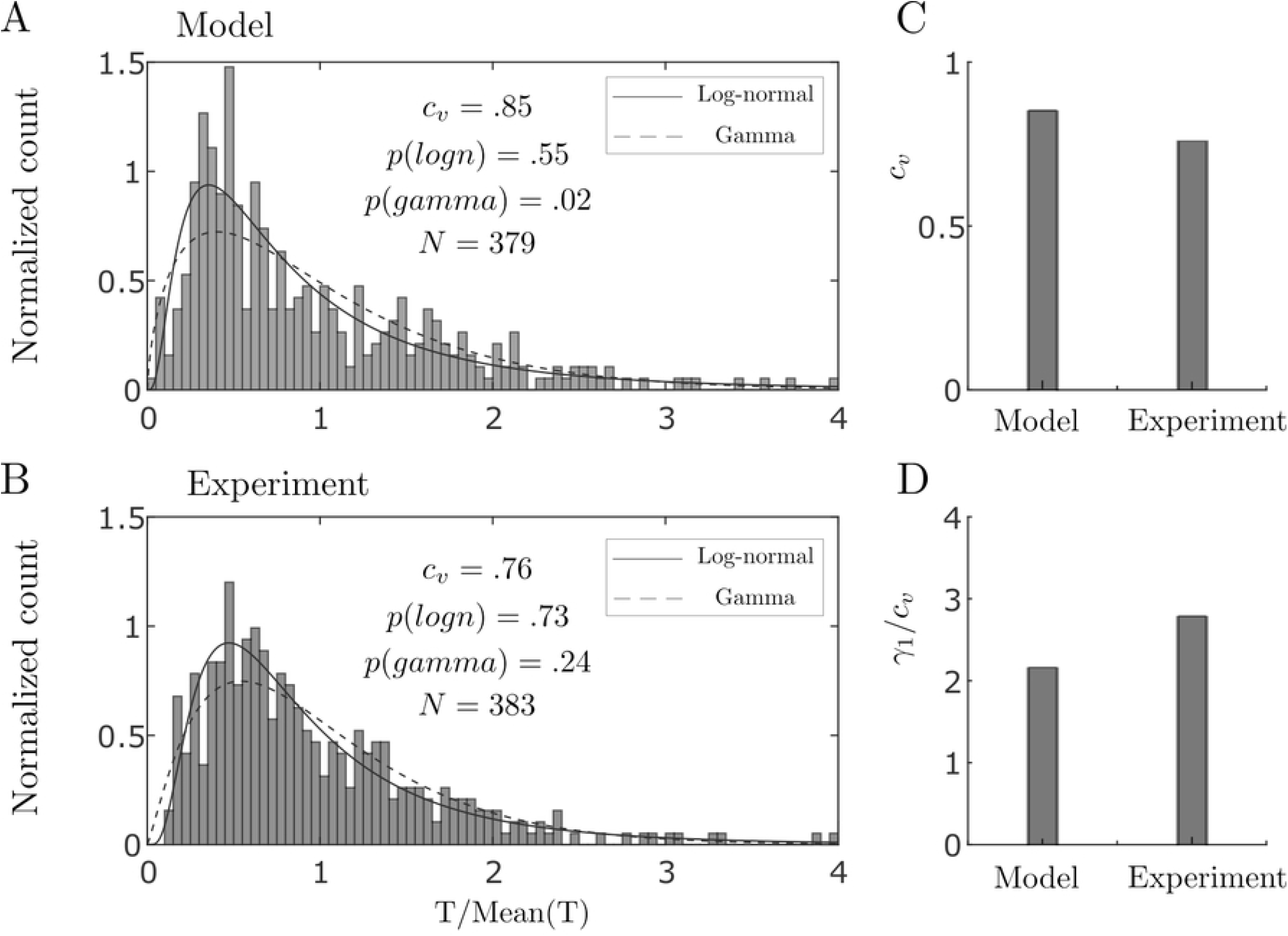
Statistics of dominance durations. **(A) Model**. Histogram of 379 durations from model simulations at Δ*I* = 2 *dB* combined across perceptual type after normalising by the mean. Solid and dashed curves show the estimated log-normal and gamma distribution, respectively. P-values are from one-way KS test. **(B) Experiment**. Histogram of normalized perceptual durations combined across participants and percept type after normalization by the mean, for experimental conditions close to equidominance (Δ*I* = 2 *dB*). **(C–D) Moment ratios. (C)** Coefficient of variation (*c*_*v*_) and **(D)** skewness divided by coefficient of variation (*γ*_1_/*c*_*v*_) computed for distributions from the model and the experiment at intensity difference Δ*I* = 2 *dB*.

To assess how well tactile rivalry model conforms to the moment distribution ratios reported in [12], the statistical characteristics are compared across the model and the experiments. As Fig 5C&D shows coefficient of variation for the distribution from the model is *c*_*v*_ = .85, and the ratio of skewness and coefficient of variation is *γ*_1_/*c*_*v*_ = 2.17, which are comparable to the corresponding values from experiment (*c*_*v*_ = .76, *γ*_1_*/c*_*v*_ = 2.79), see discussion section for more details.

## Discussion

Here, a two-stage model of tactile rivalry is introduced that encodes the temporal dynamics and features of both percepts observed in tactile rivalry experiments, and alternations between these percepts. Bifurcation analysis was used to tune model parameters for the first stage to operate within bistable or oscillatory regime. And the second stage model parameters are tuned to operate within a range where direct transitions between SIM and AM are possible. Other model parameters have been estimated through a genetic algorithm with a cost function to minimise the differences between the experimental and computational mean dominance curves with respect to intensity difference. The powerful combination of bifurcation analysis to tune certain features of the model, along with optimisation tools, allowed for the design of a model that captures many features from the experiments of tactile rivalry.

### Physiological basis of tactile rivalry model

The somatosensory cortex consists of several neighbouring, functionally distinct areas whose interconnections are complex and only partially understood. S1 and S2 are two major areas of the somatosensory cortex. The most anterior (forward) of S1’s four strips is known as area 3a, the most posterior (rear) strip is known as area 2, areas 3b and 1 lay in between (Fig 1C). The model at the first stage receives contra-lateral excitation and ipsi-lateral inhibition, assumed to be tactile nerve fibre responses to the right and left hand stimuli. Neural populations in the first stage could be located somewhere in somatosensory pathway between the brainstem nuclei (where the right and left afferent fibres cross over) and area 3b of primary somatosensory cortex (Fig 1C).

In the model, neural populations in the first stage make inter- and intra-hemispheric inhibitory connections with the neural populations in the second stage. Several studies have reported evidence for interhemispheric interactions in anterior parietal cortex during bimanual stimulation in humans [46, 47]. In area 3b of monkeys, interhemispheric interactions have been described as primarily suppressive, in that simultaneous tactile stimulation of both hands suppresses neural activity in area 3b measured on one side through optical imaging [48].

The neural populations at the second stage are plausibly located within area 3b of the primary somatosensory cortex. Neurons in this area have been characterized using linear spatial receptive fields with spatially separated excitatory and inhibitory regions [49]. In addition to an inhibitory component flanking the excitatory one, receptive fields tend to also comprise an inhibitory component co-localized with the excitatory field but delayed by 20 to 30 ms [50, 51]. This receptive field structure results in an initial excitatory drive that is followed by an inhibitory one, rendering the neuron less excitable for a period of time [49]. Considering these facts, the inclusion of intra-cortical fast excitation and delayed inhibition in the model (a delay of 5 ms is considered in the model, which is comparable to the delays observed in area 3b) can account for these excitatory and lagged inhibitory components of receptive fields. Neurons in areas 1 and 2 (higher processing areas within S1), in contrast to their counterparts in area 3b, tend to have larger and more complex receptive fields and also exhibit more complex feature selectivity [52, 53].

### Stochastic influences on perceptual switching

In neural competition models, noise and adaptation processes are two possible switching mechanisms that account for perceptual alternations [27, 54]. In consideration of the experimental constraints on the statistics of alternations (mean of the dominance durations, their coefficient of variation and correlations between successive durations), models must operate with a balance between the noise and adaptation strength [55]. In several competition models, alternations are possible over large regions of parameter space, but the experimental constraints are satisfied in only a restricted domain where precise tuning of the system’s parameters is necessary [55].

The choice of stochastic process to reproduce the characteristics of perceptual rivalry including the short-tailed skewness of reversal time distributions has recently been under investigation [56]. It has also been shown that a generalized Ehrenfest stochastic process reproduces an experimentally-observed scaling property giving consistent ratios of distribution moments across a range of parameters, and the short-tailed skewness of reversal time distributions [19]. In the present study, we used an Ornstein-Uhlenbeck process, and the results produced by simulation of the tactile rivalry model were best fit by a log-normal distribution, consistent with a recent auditory and visual bistability study involving a large number of subjects [20]. The ratio of skewness and coefficient of variation for the distribution from the model are comparable to experimental values [12].

We found significant negative correlation for successive (lag 1) perceptual durations (from SIM to AM and vice versa) in the model. For perceptual durations that were one phase apart (lag 2), the correlation was not significantly different from zero (statistically independent, see S4 Fig in S1 Appendix). These results are in contrast with tactile rivalry experiments (significant positive correlation for lag 1 and negative correlation for lag 2) [12]. For the current model to capture statistical features of multistable perception including correlations, we would need to further explore the choice of noise processes. A recent hierarchical model of binocular rivalry uses out-of-equilibrium dynamics to reproduce dependence of durations on input strength, as well as the distribution of dominance durations and correlations [23]. A further investigation on the choice of stochastic process in the present tactile rivalry model could be done without the need to change other elements of model that work well (e.g. dependence of dominance durations on input strength).

### Predictions

Experimental data was used to constrain the model, in particular, the optimisation approach presented here determined the shape of the monotonically increasing, nonlinear relationship between Δ*I* and stage 1 inputs (*D*). Equipped with this nonlinearity, the model can predict e.g. the dominance durations at values of Δ*I*. It would be interesting to explore whether the model outperforms linear extrapolation between experimental data points. Furthermore, the nonlinearity predicted from the model may underpin other computations involving detection of differences in input intensity across the left and right fingers (or more generally at other locations).

In the model, the activity in stage 1 is elevated (UP) when detected differences between left and right inputs (Fig 2A) is transiently enhanced based on intrinsic noise in the population (Fig 2B) and the current state of adaptation in the neural populations (not shown). The resulting effective enhancement of inhibition in stage 2 leads to the SIM percept. Determining brain regions that encode differences between left and right tactile inputs would shed light on how the computations in stage 1 are driven. This left-right difference could feasibly be computed after the brainstem nuclei and still be in a subcortical area. However, activity that correlates with perception (as in Fig 2C–D), and biases the encoded in stage 2 (as in Fig 2E) is more likely to be cortically based. Indeed, evidence from recordings in macaques viewing bistable stimuli show that the proportion of percept-related activity increases in higher (non primary) visual areas [57]. As discussed above, anterior parietal cortex could be involved in these computations, which could be investigated with non-invasive imaging.

Based on the available literature, we propose that stage 2 is plausibly located in area S1 3b. This is an experimentally testable prediction. For human participants, non-invasive imaging (EEG, MEG, fMRI) would not allow the spatial resolution to localise activity generated in this subdivision of S1. However, for the auditory system, the timing of activity associated with differences in perceptual interpretation can shed light on the putative origin of perceptual decisions [58]. Moreover, recent work using large-scale intracranial recordings (in patients undergoing brain surgery) offers a unique opportunity to investigate this type of prediction [59].

### Future work, Levelt’s proposition IV

In this study, we only investigated Levelt’s proposition II, which considers the relation between dominant perceptual durations and asymmetric variation of feature difference (here Δ*I*). Further experimental work needs to be done to demonstrate whether Levelt’s proposition IV also extends to tactile rivalry. This would provide an opportunity to further test and improve the model.

Further experimental work could investigate the effects of other features of the stimuli such as presentation rate (*PR*) and pulse durations (*TD*), drawing comparisons against the auditory streaming paradigm [6]. The application of bifurcation analysis to periodically forced rivalry models, originally presented in [30] and utilised here, could be used to predict how the experimental result may be affected by the variation of these stimulus parameters.

The model presented here works with the simple tactile rivalry stimuli delivered to the same locations on each hand. This model can be further developed to look at more complex stimuli such as tactile motion quartet involving four locations on the skin [8].

## Conclusion

Earlier experimental work showed that perceptual rivalry extends to the tactile domain and common characteristics of multistable phenomena also persist with vibrotactile stimuli. This study presents a mathematical model for tactile rivalry based on physiological details from the somatosensory processing pathway. The proposed model is based on plausible neural mechanisms found throughout cortex and lower brain areas, and captures the temporal and feature characteristics of perceptual interpretations for tactile rivalry. With parameter tuning model produces general characteristics of perceptual rivalry including Levelt’s proposition II, the short-tailed skewness of reversal time distributions and the ratio of distribution moments. The putative neural populations of the first stage could be located somewhere in the somatosensory pathway between the brainstem nuclei and S1 (area 3b), and the neural populations of the second stage could be located within area 3b of the primary somatosensory cortex. The presented hierarchical model is generalisable and could be adapted to account for percept formation and competition leading to alternations for perceptually bistable stimuli from visual and auditory domains.

## Supporting information

### S1 Appendix

#### Stage 1-encoding perceptual alternations

The dynamics of adapting recurrent model [35] is described in terms of the mean firing rate *ν*, and activity-driven adaptation *α* (S1 FigA):

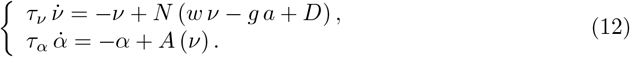

Where *w* is the strength of recurrent excitation, *g* is the strength of adaptation, *D* is the level of input, and *τ*_*ν*_ and *τ*_*α*_ are time scales of mean firing rate and adaptation, respectively. *N* (*x*) and *A*(*x*) are assumed to be sigmoidal activation function as follows;

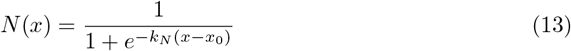

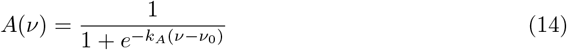

Unless otherwise specified, we use *x*_0_ = 5, *ν*_0_ = 0.5, *k*_*N*_ = 1 and *k*_*A*_ = 15 to parametrize the activation functions.

We used two of these adapting excitatory units in the fist stage of the full tactile rivalry model (Fig 1D), one for the right and one for the left side (similar equations for variables *ν*_*R*_, *α*_*R*_ for the left side, and variables *ν*_*L*_, *α*_*L*_ for the left side). Where inputs *D*_*R*_ and *D*_*L*_ to these models are the stimulus differences from the right and left as follow;

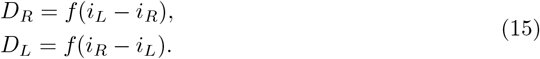

Function *f* is a nonlinearity, which will be determined later based on data-driven optimisation (see results section).

#### Stage 1-Bifurcation analysis

In the adapting recurrent model, UP/DOWN alternations are possible if there are coexisting stable states corresponding to UP and DOWN (i.e. bistability). This requires adequate strength of recurrent excitation *w*, to self-maintain the UP state under conditions of low drive. We show this first for a reduced case without adaptation dynamics (*g* = 0 in Eq (12)).

The steady states of firing rate variable *ν*, depend on the level of drive *D*, and the strength of recurrent excitation *w*. If recurrent excitation is weak (*w* = 0), the right-hand side of Eq (12) increases monotonically with *D*, and has only one solution which is a stable fixed point (S1 FigC). As recurrent excitation increases, the right-hand side of Eq (12) shows a region of bistability between a low-rate bifurcation point at weak drive and a high-rate bifurcation point at strong drive (S1 FigB). In the (*w, D*)-parameter plane, the bistable region (grey shaded area) has borders that correspond to saddle-node bifurcations (S1 FigD). UP/DOWN bistability emerges at a critical value of recurrent excitation (*w* = 4), and a critical level of drive (*D* = 3). Consequently, a first general insight of adapting recurrent models is that UP/DOWN bistability will emerge in neuronal populations with sufficiently strong recurrent excitation, during conditions of low drive.

#### Stage 2-encoding perceptual interpretation

The starting point for this model stage is a periodically driven competition network of two localised Wilson-Cowan units previously used for the encoding of different perceptual interpretations of the auditory streaming paradigm [31]. The model is described by the following system of delayed differential equations (DDEs):

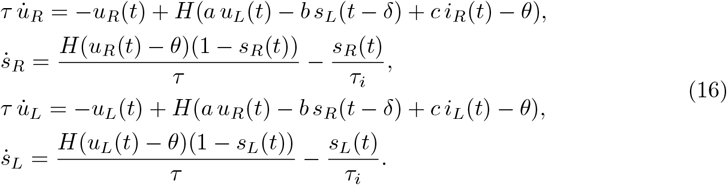

Where units *u*_*R*_ and *u*_*L*_ represent the mean firing rate of two neural populations encoding sequences of vibrotactile pulses with timescale *τ*. The Heaviside gain function *H*(*x*) with activity threshold *θ* is given by

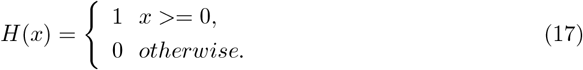

Mutual coupling through direct fast excitation has strength *a*. The delayed, slowly decaying inhibition has timescale *τ*_*i*_, strength *b* and delay *δ*. The model is driven by excitatory inputs *i*_*R*_(*t*) and *i*_*L*_(*t*) with strength *c*. The synaptic variables *s*_*R*_ and *s*_*L*_ describe the time-evolution of the inhibitory dynamics.

#### Transformation to continuous model for bifurcation analysis

As shown in [31], this system exhibits three different dynamical states relevant to perceptual interpretations of alternating stimuli (S2 FigA). The states are distinguishable by the number of threshold crossings per 2*TR* period: (1) both units (*u*_*L*_ and *u*_*R*_) cross threshold in response to both input pulses (low and high) (total of 4 crossings, corresponds to integration in auditory streaming), (2) the *R* unit crosses threshold twice and the *L* unit once (total of 3 crossings, corresponds to bistability in auditory streaming), and (3) both units cross threshold once (total of 2 crossings, corresponds to segregation in auditory streaming).

The states (1) and (3) match the percepts observed in the experiments of tactile rivalry (SIM and AM, respectively). However, the state (2) does not have any specific meaning in the tactile domain (S2 FigB). So, we are looking for a region in the parameter space that state (2) does not exist.

The model in Eq (16) has been described using a system of DDEs. To be able to perform bifurcation analysis (using Auto07p), we need to transform it to a system of ODEs. For this aim, the dynamics of direct inhibitory synapses *s*_*R*_(*t*) and *s*_*L*_(*t*) is replaced with the dynamics of an indirect synapse (to describe the dynamics of *s*_*R*_(*t* − *δ*) and *s*_*L*_(*t* − *δ*)). These indirect synapses are modelled by introducing two other synaptic variables *x*_*R*_(*t*) and *x*_*L*_(*t*) [45] with dynamics described by:

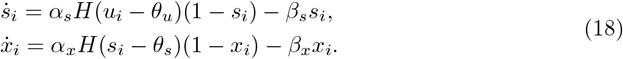

Here, *α*_*i*_ and *β*_*i*_ (*i* = *s, x*) are positive constants. These indirect synapses have the effect of introducing a delay in the synaptic action, and this delay takes place on the slow timescale. Once *u*_*i*_ crosses the threshold *θ*_*u*_, then *s*_*i*_ will activate. The activation of *x*_*i*_ must wait until *s*_*i*_ crosses the second threshold *θ*_*s*_, thus introducing a delay.

We can choose 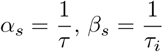, and *θ*_*u*_ = *θ*, and tune parameters *α*_*x*_, *β*_*x*_, and *θ*_*s*_, so that the dynamics of *x*_*R*_(*t*) and *x*_*L*_(*t*) can generate delays approximately equal to the fixed delay *δ* in the system of DDEs (Eq 16). Then *s*_*R*_(*t − δ*) and *s*_*L*_(*t* − *δ*) in Eq (16) can be replaced with *x*_*R*_(*t*) and *x*_*L*_(*t*), respectively. Now the dynamics of the model can be described by the following system of ODEs:

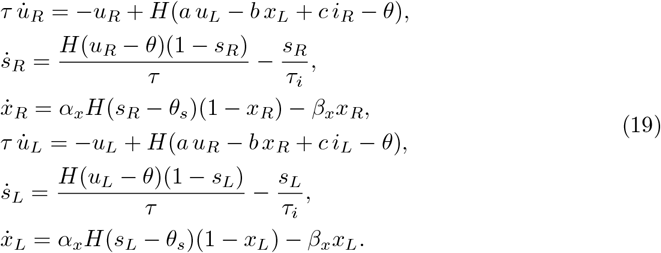

#### Stage 2-Bifurcation analysis

Bifurcation analysis of the second stage with respect to intensity difference Δ*I* is shown in S3 Fig. The left and the right units receive inputs *i*_*L*_ and *i*_*R*_, which are antiphase sequences of high and low amplitude pulses (Δ*I* below the full amplitude pulse) with duration *TD* (S3 FigA). There are reciprocal excitatory (with strength *a*) and inhibitory (with strength *b*) connections between the two units (S3 FigB). Population activities *u*_*L*_ and *u*_*R*_ can encode the SIM percept, when *u*_*L*_ and *u*_*R*_ have a full response to both pulses (S3 FigC). The AM percept is encoded when each unit only responds to the full amplitude input pulse and does not respond to the low amplitude input pulse (S3 FigD). One parameter bifurcation diagrams are shown at three different values of mutual excitation *a*, with other parameters fixed (*b* = 2.8, *c* = 5.5). A blue curve shows the branch of periodic orbits, for which there is a sharp transition between periodic responses encoding the SIM and AM percepts (top panel in S3 FigE). The boundary between these two dynamical behaviours moves to the right with higher values of excitation (middle panel), and to the left with lower values of excitation (bottom panel). When coupled to the first stage, the effective value of *a, a*_eff_ = *a* − *dν* creates a hysteresis loop between these two cases (Fig 3B).

#### Full tactile rivalry model

To form the full tactile rivalry model, two units of adapting recurrent model encoding alternations at the first stage Eq (12–15) is incorporated with the model encoding percepts at the second stage Eq (19). Units *ν*_*R*_ and *ν*_*L*_ in the first stage make inter- and intra-hemispheric inhibitory connections with strength *d* to the units *u*_*R*_ and *u*_*L*_ in the second stage (Fig 1D).

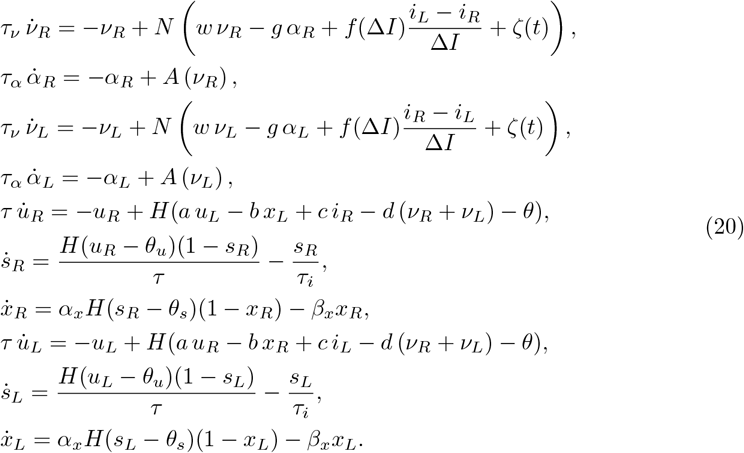

#### Simplified tactile rivalry model

As the inputs (*i*_*L*_ − *i*_*R*_) and (*i*_*R*_ − *i*_*L*_) to the full tactile rivalry model are antiphase and there is also symmetry in the first stage of the model, this model can be simplified, and units *ν*_*R*_ and *ν*_*L*_ can be replaced by one adapting recurrent model with variables *ν* and *α* and input *D* = *f* (Δ*I*) (Fig 3A). Where Δ*I* is a positive constant, and *f* is a nonlinearity to be determined by data-driven optimisation. So the simplified tactile rivalry model is described by

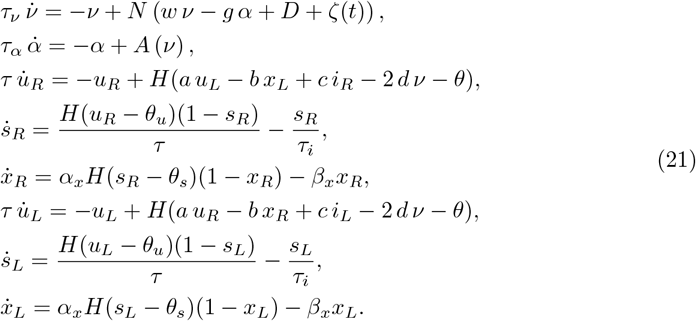

**S1 Fig.**
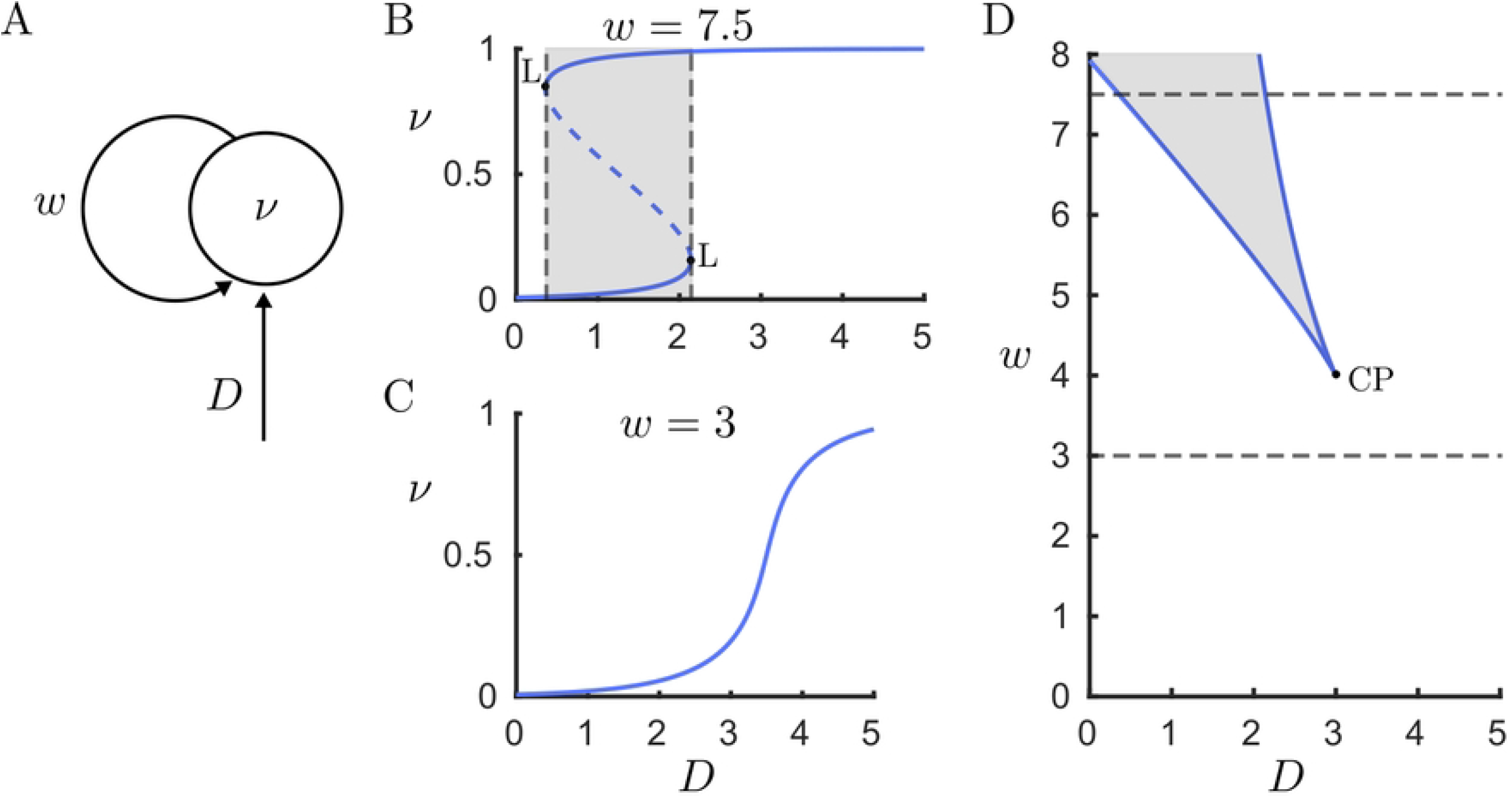
Bifurcation analysis of the adapting recurrent model (without adaptation *g* = 0). Mechanism of bistability with sufficient levels of recurrent excitation and input (high levels of recurrent excitation and low levels of input). **(A) Adapting recurrent model**. Population firing rate *ν* driven by *D* and recurrent excitation with strength *w*. **(B & C) One parameter bifurcation diagram**. Population rate steady state as a function of input *D*, for a population with low (*w* = 3), and high (*w* = 7.5) level of recurrent excitation. Stable and unstable fixed points are indicated with solid and dashed curves, respectively. **(D) Two parameter bifurcation diagram**. *W − D* parameter space, bistable region (grey shaded area) surrounded by two fold bifurcations (L) which merge together and disappear through a cusp bifurcation (CP).

**S2 Fig.**
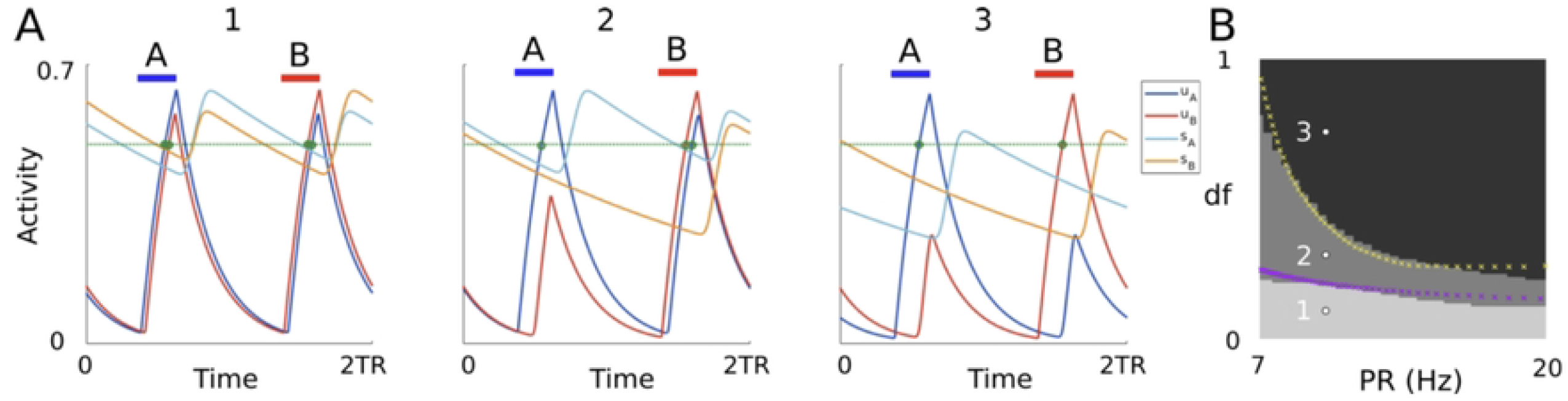
**(A)** Time histories of the 2TR-periodic states in the system defined in Eq (16). Units’ threshold crossings are shown by green dots. **(B)** The total number of threshold crossings for both units is shown in greyscale for simulated trajectories at varying *PR* and *df* (black = 2, gray =3, lightest gray = 4 crossings). Parameters *PR* and *df* in panel (A) are shown by white dots in panel (B). The remaining parameters are *a* = 2, *b* = 2.8, *c* = 5.5, *δ* = 0.015, *TD* = 0.022, *τ*_*i*_ = 0.25, *τ* = 0.025, *θ* = 0.5. Adapted from [31].

**S3 Fig.**
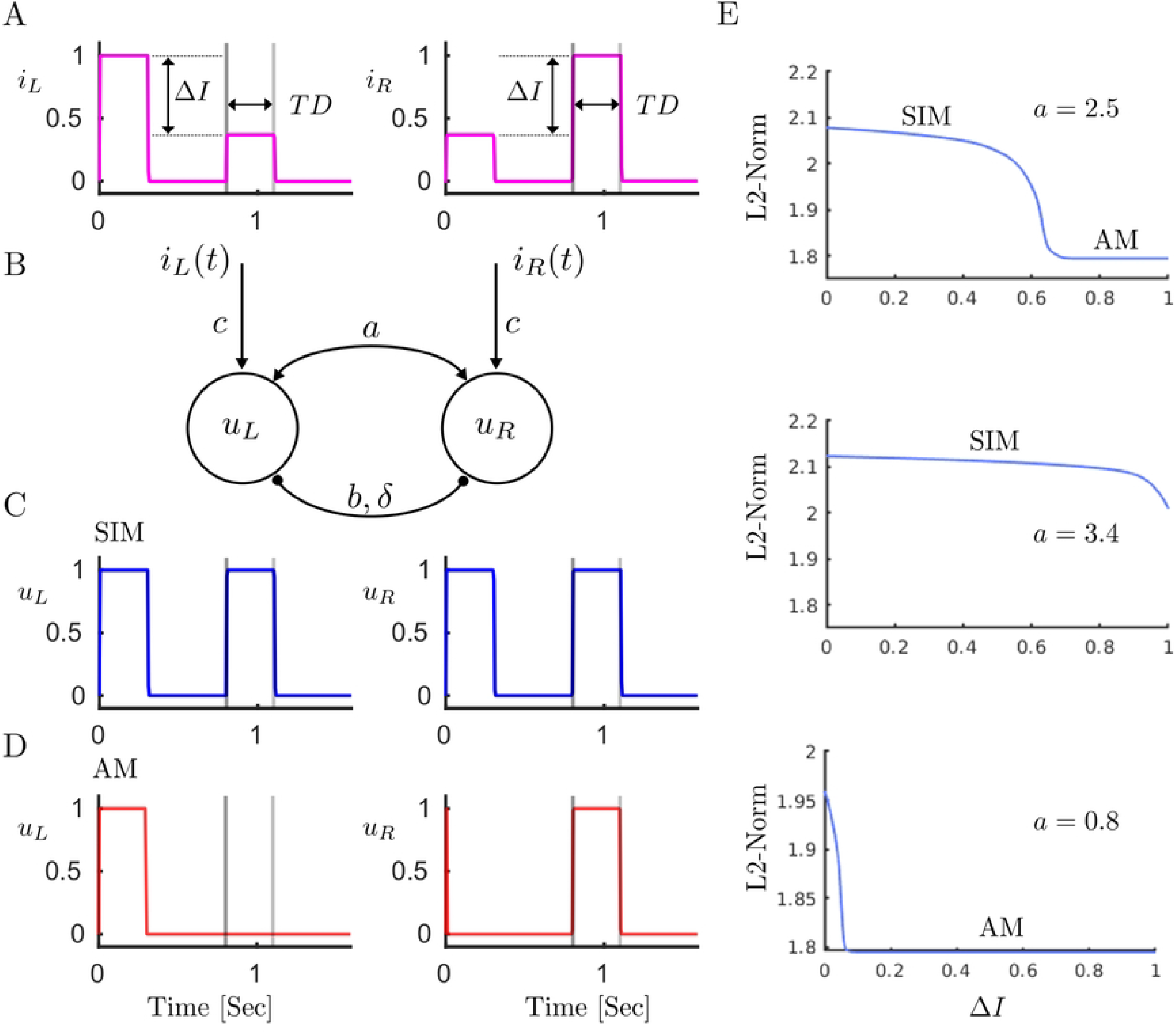
Mechanism of encoding percepts. **(A) Stimuli**. One period (length 2*TR* = 1.6 *s*) of the stimuli to the left and right units consists of one full amplitude pulse (length *TD* = 0.3 *s*), and one pulse that its amplitude is Δ*I* below the intensity of the full amplitude pulse on a logarithmic scale (*dB*). **(B) Model encoding percepts**. Schematic of the model consisting of mutual fast excitation with strengths *a* and delayed inhibition with strength *b*. Inhibition is delayed of the amount *D*. **(C) SIM percept**. Model can encode percept SIM, when *u*_*L*_ and *u*_*R*_ have full respond to both pulses. **(D) AM percept** is encoded when they only respond to the full amplitude pulse and no response to the low intensity pulse. **(E) Bifurcation analysis with respect to intensity difference** Δ*I*. One parameter bifurcation diagram sketched at three different values of *a*. Blue curve shows the branch of periodic orbit, in which there is a sharp transition between periodic respond encodes SIM and AM percepts (top panel). For higher values of excitation (middle panel) this boundary moves to the right, and for lower values of excitation (bottom panel) moves to the left.

**S4 Fig.**
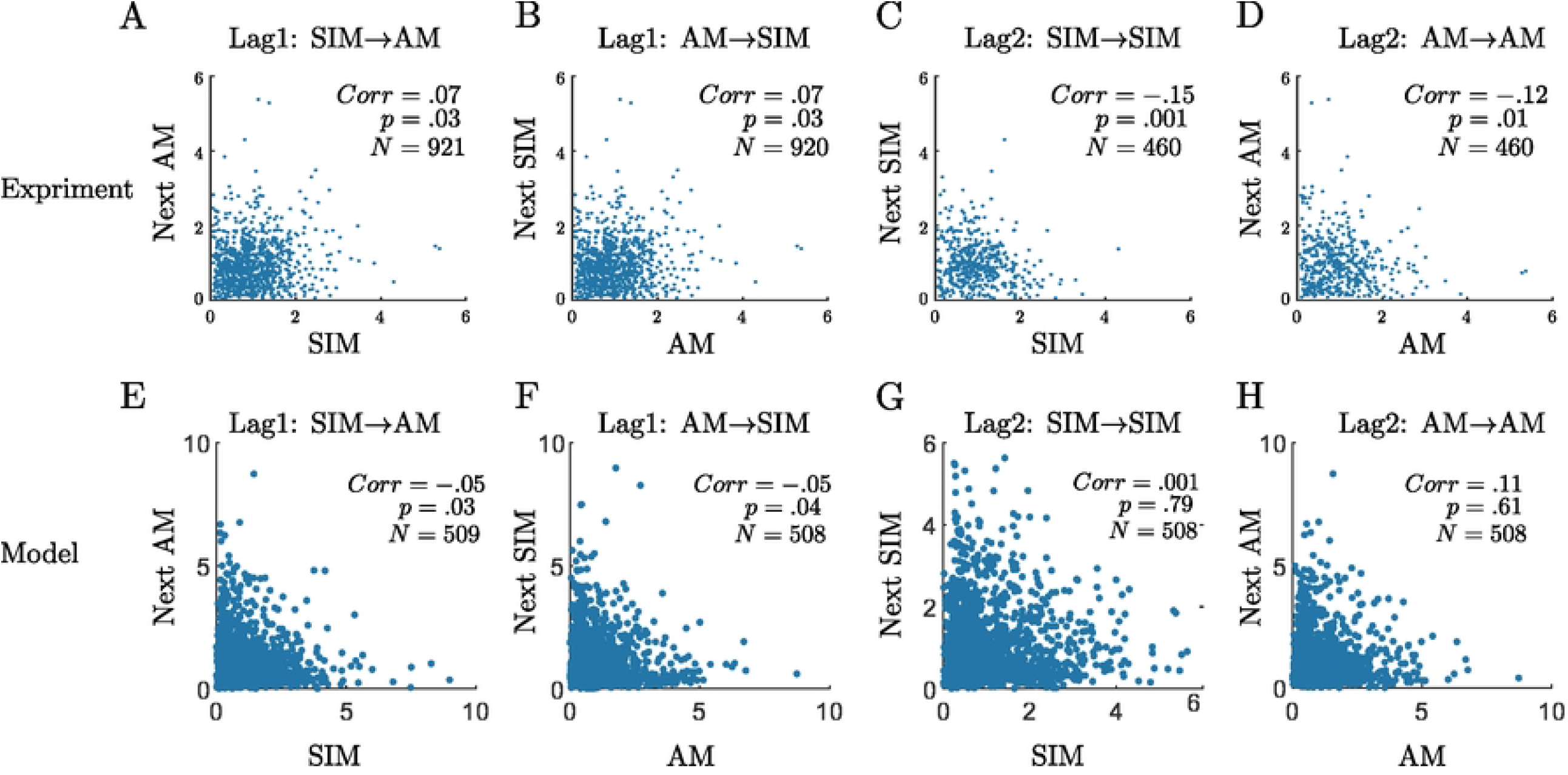
**(A-D)** Scatter plots of normalized durations from experiment. The correlation coefficient (*corr*) between perceptual phases for each scatter is indicated in each panel with the corresponding *p*-value and number of pairs (*p* and *N* respectively). **(E-H)** Scatter plots of normalized durations from model. The transition types are marked above each histogram **(A&E)** AM following SIM. **(B&F)** SIM following AM. **(C&G)** SIM. **(D&H)** AM.

